# Strawberry additive increases nicotine vapor sampling and systemic exposure but does not enhance Pavlovian-based nicotine reward in mice

**DOI:** 10.1101/2022.10.19.512743

**Authors:** Theresa Patten, Natalie L. Johnson, Jessica K. Shaw, Amanda M. Dossat, Allison Dreier, Bruce A. Kimball, Daniel W. Wesson, Mariella De Biasi

## Abstract

Nicotine is an addictive drug whose popularity has recently increased, particularly among adolescents, due to the availability of electronic nicotine devices (*i*.*e*., “vaping”) and nicotine e-liquids containing additives with rich chemosensory properties. Some efforts to understand the role of these additives in nicotine reward suggest that they increase nicotine reward and reinforcement, but the sensory contributions of additives, especially in their vapor forms, are largely untested. Here, to better understand how a fruit-flavored (*i*.*e*., strawberry) additive influences nicotine reward and aversion, we used a conditioned place preference (CPP) procedure in which nicotine and a strawberry additive were delivered as a vapor to male and female adolescent mice. We found that nicotine vapor alone can lead to dose-dependent CPP when using a biased design. The strawberry additive did not produce CPP on its own, and we did not observe an effect of the strawberry additive on nicotine vapor-induced reward. Nevertheless, mice exposed to nicotine + strawberry additive vapor had higher plasma cotinine concentrations, which did not appear to reflect altered nicotine metabolism. Instead, by directly measuring vapor sampling through respiration monitoring, we uncovered an increase in the amount of sniffing toward strawberry-containing nicotine vapor compared to nicotine vapor alone. Together these data indicate that chemosensory-rich e-liquid additives may enhance the perceived sensory profile of nicotine vapors rather than the reward value *per se*, which leads to overall increased nicotine exposure.

**Significance Statement:** With the rise in popularity of flavored e-cigarette products, many have considered the possibility that flavor volatiles will enhance nicotine reward; however, the possibility that flavor additives have chemosensory properties that can affect nicotine intake has been largely overlooked. Here, by delivering nicotine to adolescent mice as a vapor we were able to consider both possibilities. We found that mice had increased sniffing intensity and nicotine exposure when vapors contained a strawberry additive, despite the fact that the same additive was unable to enhance Pavlovian nicotine reward using a CPP paradigm. This research highlights the importance of considering the chemosensory properties of e-cigarette additives as a mechanism for their effect on nicotine use.

## Introduction

The number of adolescent nicotine users in the US increased in 2019 for the first time in decades, an effect driven by a rise in e-cigarette use (Centers for Disease Control and Prevention, 2019). Chemosensory-rich additives in e-cigarette liquids, also known as “flavors”, promote nicotine use in humans, possibly due to reduced perceptions of harm (Chaffee et al., 2018; Ford et al., 2016; Pepper et al., 2016). In addition, sweet additives such as those engineered to be perceived as fruit- or candy-like are attractive to adolescents and young adults who have a higher preference for sweetness and sensory cues, including odors, associated with sweetness (Desor and Beauchamp, 1987; Hoffman et al., 2016; Zandstra and de Graaf, 1998).

Chronic nicotine exposure via combustible cigarettes is associated with serious health conditions and is the leading cause of preventable death in the developed world (Creamer et al., 2019). Given that e-cigarette use is associated with a progression to combustible cigarette use, there is concern that e-cigarettes and their attractive additives have ushered in another generation of individuals who will come to struggle with physical and psychological health complications, such as an increased likelihood of drug dependence and increased risk of cognitive deficits (Goriounova and Mansvelder, 2012; Soneji et al., 2017).

E-cigarette additives with significant chemosensory properties are widely hypothesized to increase nicotine reward (Audrain-McGovern et al., 2016; Goldenson et al., 2016; Kim et al., 2016; A. M. Leventhal et al., 2019; Patten and De Biasi, 2020), as sensory perception plays a role in nicotine reward, use, and craving. Blocking the airway sensory experience of smoking with a local anesthetic reduces smoking satisfaction and sensory stimulation with other irritants (*e*.*g*., citric acid) can reduce cigarette cravings (Levin et al., 1990; Rose et al., 1984). E-cigarettes with fruity chemical additives are perceived as sweet, and flavor-enhanced sweetness and “liking” are often correlated in clinical reports of subjective reward (Goldenson et al., 2016; Kim et al., 2016; A. M. Leventhal et al., 2019). “Sweetness” is rewarding and reinforcing and can support both CPP and self-administration in rodents (Bacon et al., 1962; Dufour and Arnold, 1966). It is possible that the sweet reward contributed by e-cigarette additives synergizes with the drug reward contributed by nicotine to enhance overall reward. Users also report that flavored additives mask the harshness of the cigarette taste and that the addition of a fruit additive to nicotine-containing e-cigarettes suppresses unappealing sensations (Chen et al., 2019; Kim et al., 2016; A. Leventhal et al., 2019).

We investigated the role of a strawberry flavored e-cigarette liquid on nicotine reward using a conditioned place preference (CPP) paradigm in which nicotine was delivered to adolescent mice as a vapor. This route of drug delivery during Pavlovian conditioning allowed us to simultaneously consider the pharmacological and sensory components of e-cigarette reward and aversion. We next monitored the inhalation patterns of mice as they engaged with nicotine vapor with or without a strawberry additive. Together, our results begin to unveil novel sensory influences on e-cigarette use, preference, and reinforcement.

## Materials and Methods

### Animals

Adolescent male (n=191) and female (n=170) C57BL/6J mice between postnatal days (PND) 28 and 49, corresponding roughly to humans aged 12-18 year old(Yuan et al., 2015), were used in the CPP and blood collection experiments (Figures 1-5). They were housed at [Author University] with a reverse 12hr light/dark cycle (lights off at 10 A.M.) in a temperature-controlled room (24±2°C, relative humidity 55±10%). All behavioral testing and e-cigarette vapor exposures occurred in the dark phase of the light cycle. Animals were either purchased from Jackson Labs or bred in-house. Those bred in-house were the offspring of 20+ different breeding pairs and distributed throughout treatment groups to ensure adequate genetic diversity. Similarly, adolescent male (n=14) and female (n=11) C57BL/6J mice between PND 28 and 49 were used in the plethysmography experiments (Figures 4-5) and were housed at [Author University] in comparable conditions. All mice in this study had *ad libitum* access to food and water. All procedures were approved by the appropriate Institutional Animal Care and Use Committee and followed the guidelines for animal intramural research from the National Institutes of Health.

**Figure 1.**
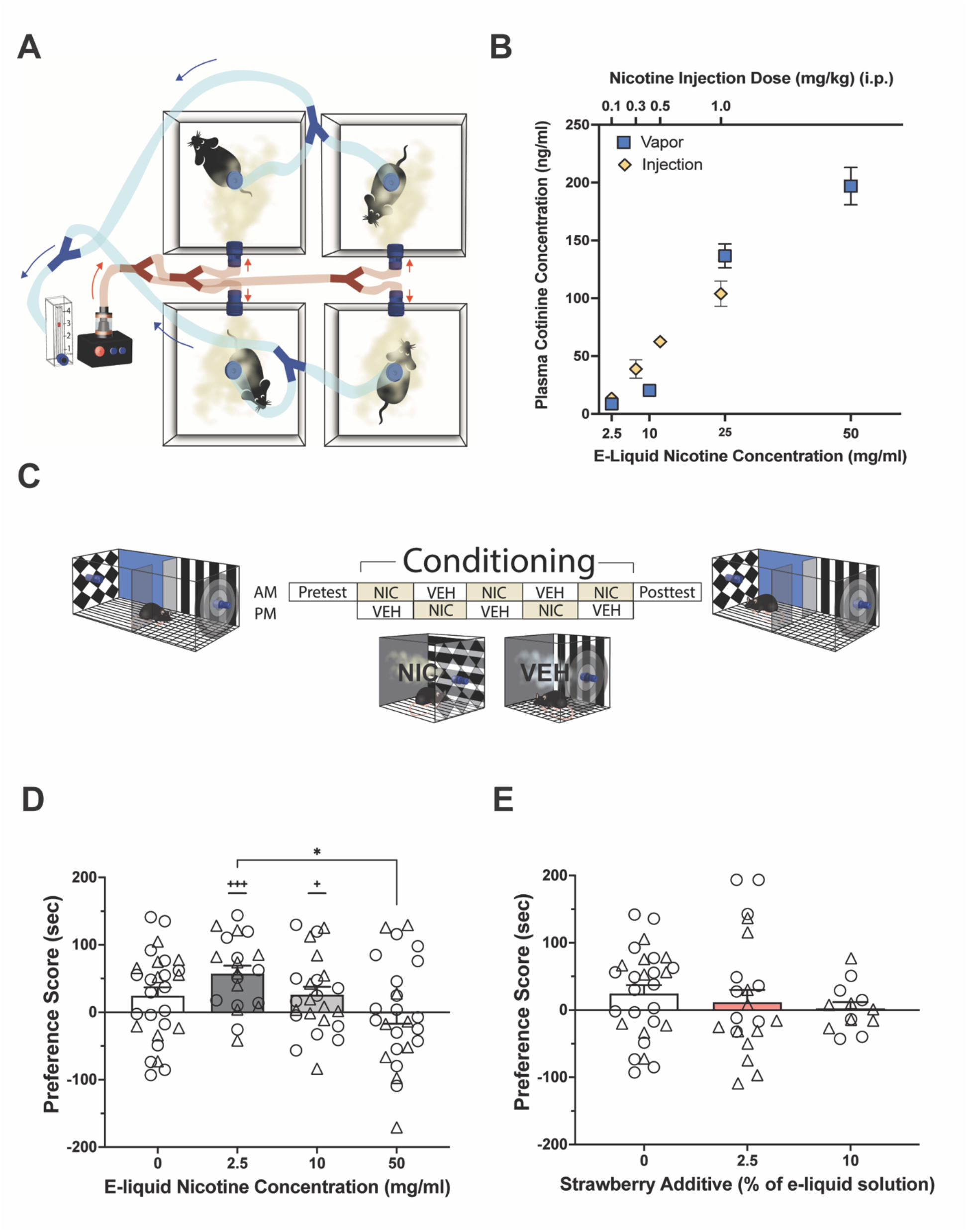
Nicotine delivery via e-cigarette vapor results in measurable plasma cotinine concentrations, and the rewarding properties e-cigarette vapors can be evaluating using conditioned place preference. (A) A vapor exposure set-up allows for the simultaneous exposure of four mice with e-cigarette vapors. (B) Scatter plots of plasma cotinine concentrations detectable in adolescent mice (mean ± sem) 30 minutes after nicotine exposure via an intraperitoneal injection (yellow diamonds, top x-axis, n=7,4,7,5) or e-cigarette vapor exposure as depicted in panel A (blue boxes, bottom x-axis, n=27, 28, 4, 46). (C) Schematic of the vapor conditioned place preference paradigm in which mice receive experimental vapor (*e*.*g*., nicotine) once per day for 5 days in the CS+ compartment and vehicle once per day for 5 days in the CS-compartment. (D) Scatter plots show preference scores ((time in CS+ during posttest) – (time in CS+ during pretest)) of adolescent mice conditioned to nicotine vapor of various concentrations using a CPP-biased method (n=28,20,24,25). (E) Scatter plots show preference scores of individual adolescent mice conditioned to nicotine vapor of various concentrations using a CPA-biased method (n=29,21,23,21). In panels D & E, bars show the mean ± sem and the vehicle data is the same in both panels. One sample t-tests on change in time spent in CS+ (i.e. vs 0 second change in time spent in CS+): +p<0.05, +++p<0.001. One-way ANOVA and multiple comparisons: *p<0.05: relative to comparison group. For all charts, circles represent individual male mice and triangles represent female mice.

### E-liquid and Nicotine Preparations

E-liquid materials [Vegetable Glycerin (VG), Propylene Glycol (PG), NicSelect Nicotine (free-base; 100 mg/ml in VG), and Strawberry Flavor Concentrate (in PG)] were purchased from Liquid Barn™ (Simi Valley, CA). E-liquids were mixed in the lab to the desired concentration of nicotine and strawberry additive while maintaining a 50/50 ratio of VG/PG. E-liquid was kept in the dark and made fresh every three days to prevent nicotine degradation. pH was not adjusted between solutions. For intraperitoneal (i.p.) nicotine injections, nicotine tartrate salt (Sigma Aldrich, St. Louis, MO) was dissolved in Phosphate Buffered Saline (PBS) to desired concentrations (0.03-0.1 mg/ml freebase nicotine). When the additive was added to PBS for injections, a concentrated nicotine stock solution was used to make both the nicotine only and nicotine+additive injection solutions. The final injection solution was diluted with either PBS alone or PBS containing 5% strawberry additive (Liquid Barn™, Simi Valley, CA) to ensure that nicotine concentration in the injection solutions was identical.

### E-cigarette Vapor Delivery

E-liquid was vaporized using a Vapor Generator Controller (La Jolla Research Inc, La Jolla, CA) in conjunction with a SMOK Baby Beast Brother e-cigarette tank (SMOK, Shenzhen, China). The Vapor Generator Controller was operated at 75.0W using the system’s pre-set ‘nicotine’ settings. This machine has been previously validated for nicotine vapor delivery (Cooper et al., 2021; Montanari et al., 2020). The vapor ‘puff’ from a single e-cigarette is diverted into four air-tight exposure apparatuses (dimensions=5.25” x 5.25” x 5.625”) and is quickly replaced by clean room air via a vacuum (flow rate=3.0 L/min) (for schematic, see Figure 1A). We delivered a 1-s puff every 90 s for a total of 25 mins, resulting in a vapor period which, in pilot experiments, we found was sufficient to allow delivery of pharmacologically relevant doses of nicotine while delivering vapor in a puff-like pattern. This delivery pattern also provided sufficient time for perception of the cues and association of cues with the presence (or absence) of nicotine. The 25 min vapor delivery period is followed by a 5 min ‘wash out’ to protect experimenters from vapor inhalation. Therefore, mice are placed individually into one of the four exposure chambers for a total of 30 mins per exposure. When vapor exposures were part of a conditioned place preference or aversion experiment, tactile and visual cues were added to the floor and to the outside of the clear plexiglass chamber (see Figure 1C). Tactile cues comprised either rigid embroidery mesh in a grid pattern or flexible corrugated floors.

### Cotinine Quantification

To measure plasma cotinine (the primary metabolite of nicotine) concentrations in mice following nicotine exposures, trunk blood was collected 30 mins following nicotine treatment. 30 mins post-injection has been previously shown to be the approximate time at which the C_max_ of plasma cotinine is achieved in C57/BL6 mice (Siu and Tyndale, 2007), and pilot studies in our lab indicate that this is also true using our vapor delivery system (unpublished observations). After blood collection and plasma separation, cotinine was quantified using a Mouse/Rat Cotinine ELISA kit according to package instructions (Calbiotech Inc., El Cajon, CA). Samples were diluted with purified, Milli-Q water when necessary to achieve a concentration of cotinine within the range of the standard curve that allowed for the most accurate cotinine quantification (10-50ng/ml), and a pipetting variability of <15% was required for inclusion of data. Samples that did not meet these requirements were re-analyzed. Cotinine values in diluted samples were calculated as: cotinine=cotinine_dilution_*dilution factor.

### Analyses of strawberry additive volatiles and e-liquid nicotine concentration

To identify major volatiles in the Strawberry Flavor Concentrate (Liquid Barn™, Simi Valley, CA), headspace gas chromatography/mass spectrometry (GC/MS) analyses were conducted with a HT3 dynamic headspace analyzer (Teledyne Tekmar, Mason, OH, USA) outfitted with Supelco Trap K Vocarb 3000 thermal desorption trap (Sigma-Aldrich Co., St. Louis, MO, USA). The GC/MS (Thermo Scientific Trace Ultra) was equipped with a single quadrupole mass spectrometer (Thermo Scientific, Waltham, MA, USA) and a 30 m x 0.25 mm id Stabiliwax®-DA fused-silica capillary column (Restek, Bellefonte, PA, USA). E-liquid additive solutions (0.1% in water) were subjected to dynamic headspace GC/MS analysis by placing 50 µL in a sealed 20mL headspace vial. The vial was maintained at 30°C, swept with helium for 10min (flow rate of 75mL/min), and the volatiles were collected on the thermal desorption trap. Trap contents were desorbed at 260°C directly into the GC/MS using a split injection. The GC oven program had an initial temperature of 40°C (held for 3.0min) followed by a ramp of 7.0°C/min to a final temperature of 230°C (held for 6.0min). The MS was used in scan mode from 33 to 400 m/z with a three minute solvent delay. Mass spectral peak identifications were assigned based on the library search of the NIST Standard Reference Database.

### Nicotine Vapor Conditioned Place Preference

The CPP procedure is divided into three phases: pre-test, conditioning, and post-test (see Figure 1C). All phases of testing occurred during the dark-cycle in either dim (1-2 Lux) or red light to prevent circadian rhythm disruption. During the pre-test, mice were able to freely explore a testing chamber with two rooms containing identical cues (*e*.*g*., tactile, visual) and dimensions as the vapor exposure chamber (described in ‘E-cigarette Vapor Delivery’ section) for 15 minutes. One chamber had the following cues: walls: black and white “bulls-eye” symbol, solid white, vertical black and white stripes, and solid black; flooring: rigid white embroidery mesh in a grid pattern. The opposing chamber had the following cues: walls: black and white argyle pattern, solid blue, horizontal black and white stripes, and solid black; flooring: flexible corrugated floors. CPP was conducted using a biased design (i.e., nicotine was assigned to the least preferred chamber). During the 5-day conditioning phase, mice received one CS+ (conditioned rewarded stimulus) and one CS-(conditioned unrewarded stimulus) conditioning session each day, in which they were placed in an exposure chamber individually with appropriate cues and received either control (50/50 VG/PG blend; vehicle) vapor, strawberry vapor, or nicotine ± strawberry flavored vapor. A control group received vehicle vapor in both compartments. AM and PM exposure sessions occurred approximately four hours apart. Following conditioning, mice were placed back into the testing chamber and allowed to freely explore both chambers for 15 minutes in a drug-free state. To avoid extreme bias skewing the results, mice showing strong innate preference (*i*.*e*., >65%) towards either compartment during the initial pretest were excluded.

Videos of both the pre-test and post-test were recorded and an experimenter blinded to the test stage (pre- or post-test) and to the treatment group scored time spent in each compartment, as well as the time spent climbing in the center region (unfocused). Preference scores were determined by comparing the time spent in the treatment-paired compartment (CS+) before and after conditioning with e-cigarette vapors (i.e., preference score = time in CS+ after conditioning – time in CS+ before conditioning), as this allows for a detection in the change in preference (for review, see McKendrick and Graziane, 2020).

### Cotinine Category Determination

As the inclusion of the strawberry vapor increased nicotine intake (measured by cotinine plasma levels), thereby altering the dose of nicotine received, we sought to compare mice based on the actual level of nicotine exposure. ‘Cotinine Categories’ were objectively defined according to plasma cotinine concentrations following injection of nicotine doses previously shown to be “subthreshold” (0.1 mg/kg), “rewarding” (0.3-0.5 mg/kg) and “aversive” (1.0 mg/kg) in rodents (Fudala et al., 1985; Jorenby et al., 1990; Torres et al., 2008). In our injection studies, plasma cotinine concentrations following a 0.3 mg/kg were 47.75 ± 19.05 ng/ml (mean ± s.d.). We therefore set the lower threshold of the cotinine concentration defined as ‘rewarding’ at 28.7 ng/ml, i.e., one standard deviation below the average cotinine concentration of a “rewarding” 0.3 mg/kg nicotine injection. The upper threshold of ‘rewarding’ plasma cotinine concentrations was set to 91.35 ng/ml, one standard deviation above the average cotinine concentration following a “rewarding” 0.5 mg/kg i.p. nicotine injection, thereby ensuring that we captured a large range of cotinine values associated with a rewarding dose of nicotine. ‘Subthreshold’ cotinine concentrations were determined to be anything below the rewarding threshold (i.e. <28.7ng/ml), and ‘aversive’ cotinine concentrations were defined as cotinine concentrations above the rewarding threshold (< 91.35 ng/ml).

### Plethysmography

E-liquids were prepared as above, and their volatile odors were presented to the mice through an air dilution olfactometer (Figure 4A). During experimentation, all stimuli were contained in 40mL glass headspace vials at room temperature. Odors mixed with medical-grade nitrogen (Airgas) passed through headspace vials at a flow rate of 50ml/min, following which they were mixed with a carrier stream of clean filtered room air (1L/min; Tetra Whisper, Melle, Germany) before being introduced into the plethysmograph. All odors were handled within independent tubing to prevent cross-contamination.

To monitor respiration of mice in response to e-cigarette odors, we employed whole-body plethysmography and a computer-controlled olfactometer. The unrestrained whole-body plethysmograph (Data Sciences International, St. Paul, MN) allowed for detection of respiratory transients as mice freely explored the chamber. Transients were detected using a flow transducer (Data Sciences International) and digitized at 300Hz (0.1-12Hz band pass; Synapse software, Tucker Davis Technologies, Alachua, FL), following a 500x gain amplification (Cyngus Technology Inc, Southport, NC). The olfactometer allowed for precise control of odor delivery into the plethysmograph chamber (Figure 4B). User-initiated openings of valves (Parker Hannifin, Cleveland, OH) via a computer and digital relay (LabJack, Lakewood, CO) allowed for flow of vapors into the bottom of the plethysmograph for perception by the mouse. Following each stimulus trial, odor vaporized air was passively cleared from the plethysmograph through an exhaust outlet at the chamber’s ceiling since the filtered room air is continuously delivered.

Mice were acclimated to the plethysmograph for two days prior to testing. During acclimation, mice were placed in the plethysmograph for 30 mins and vehicle solution (50/50 VG/PG) was presented for 10 s each with a 60 s inter-trial interval (ITI). The goal of this was to allow the mice to acclimate to handling, the chamber, and other possible non-olfactory cues (pressure changes, sounds) associated with valve opening and air flow. The day following this acclimation, mice were exposed to e-cigarette odors. On this day, first VG/PG was presented for 10 s with a 60 s ITI, to further acclimate the mouse to vapor delivery, followed by five trials of pseudorandom presentations of 2.5% strawberry, 10 mg/ml nicotine ± 2.5% strawberry, and 50 mg/ml ± 2.5% strawberry. Each odor was presented for 10 s with a 90 s ITI.

Inhalation peaks were detected offline in Spike2 (Cambridge Electronic Design, Cambridge, UK). First, the respiratory data were filtered (0.1-12Hz band pass). Second, the maximum point of each respiratory cycle was identified, and instantaneous frequency was then calculated based off of the duration between one peak and its preceding peak. Data were down-sampled to 50Hz for subsequent analyses. Although the odor valves were open for 10 s, the first second was excluded from analysis to account for the latency of odor delivery to the plethysmograph (Figure 5B) and unavoidable pressure artifacts (e.g., Figure 5C). We calculated the average instantaneous frequency over the remaining nine seconds of each odor presentation to quantify odor-evoked sniffing. Amplitude was calculated based on the root mean square of the respiratory signal.

### Statistical Analyses

All data sets were tested for normality using the D’Agostino Pearson Test prior to selecting appropriate methods of analysis (nonparametric vs. parametric). Nonparametric data (i.e. data that were not normally distributed) were analyzed using Mann-Whitney and Wilcoxon tests, and the Scheier-Ray-Hare test—an extension of the Kruskal-Wallis that allows for a factorial design— was used to detect interactions. Parametric tests employed in this study include t-tests, fixed effects one- and two-way ANOVAs. GraphPad Prism (San Diego, CA) was used for all data analyses except the Scheirer-Ray-Hare test, which was conducted using the Real Statistics Add-In for Microsoft Excel. In all figures, asterisks (*) indicate a significant difference from a comparison group, a plus sign (+) indicates a significant difference from 0 (0 s change from pretest; no change in preference from baseline). Mice were excluded from CPP experiments in one of two instances: 20% of screened mice (n=39/194) showed a strong initial preference (>65%) for a particular compartment during the pre-test, and two mice climbed out of the apparatus during the pre- or post-test. Sufficient numbers of mice of each sex were used to detect sex effects in behavioral tests (≈7) and in cotinine quantification experiments (≈5) when possible. Sex differences were always investigated using ANOVAs; data were collapsed across sex when a significant difference between sexes was not identified. Final sample sizes are included in figure legends.

## Results

### Nicotine vapor delivers nicotine at behaviorally-relevant concentrations

While it is known that systemic nicotine injections produce CPP/CPA (*e*.*g*., (Fudala et al., 1985; Jorenby et al., 1990; Torres et al., 2008), we wanted to assess the ability of nicotine vapor to produce these behaviors. We first validated a vapor delivery system that allowed the delivery of vapor to four chambers simultaneously (Figure 1A). To ensure appreciable nicotine delivery using this method, we measured plasma cotinine concentrations in adolescent mice (postnatal day 28-49) 30 mins after nicotine vapor exposure at three e-liquid nicotine concentrations (2.5, 10, 50 mg/ml). The resulting plasma cotinine concentrations were compared to those resulting from i.p. nicotine injections, a traditional route of nicotine delivery in preclinical research (Figure 1B). Nicotine injections consisted of a dose typically below the threshold needed to produce CPP (0.1 mg/kg nicotine), two doses previously shown to produce CPP (0.3, 0.5 mg/kg nicotine), and a dose capable of producing CPA in mice (1.0 mg/kg nicotine) (Fudala et al., 1985; Jorenby et al., 1990; Torres et al., 2008). Nicotine delivery via both routes of administration produced linearly increasing concentrations of plasma cotinine 30 mins post-nicotine treatment in adolescent mice. Plasma cotinine concentrations achieved following vapor delivery were within a range known to be behaviorally relevant, as determined by cotinine concentrations measured in adolescent mice (Figure 1B) at nicotine doses previously shown to affect behavior following injection (Fudala et al., 1985; Jorenby et al., 1990; Torres et al., 2008).

### Nicotine vapor produces CPP in adolescent mice

Next, we adapted our vapor delivery system for use in a novel vapor CPP protocol (Figure 1C; see methods). Briefly, during a pre-test, we recorded baseline preferences for both compartments. Mice were then conditioned for five days with e-cigarette vapors, and a preference score was calculated comparing the time in the treatment-paired compartment (CS+) after conditioning to the time in the CS+ during the pre-test. This widely used approach to preference score calculation allows us to detect changes in preference following conditioning (McKendrick and Graziane, 2020). A control group received vaporized vehicle containing equal parts vegetable glycerin (VG) and propylene glycol (PG), industry-standard e-cigarette solvents, in both compartments.

Nicotine vapor produced significant CPP. Mice increased their time spent in the CS+ compartment relative to baseline at the two lower nicotine concentrations tested (2.5 and 10 mg/ml unflavored nicotine e-liquid) but did not at 50 mg/ml nicotine in the e-liquid: one-sample t-tests, vs 0s: (0 mg/ml nicotine) = t(27)=2.01, p=0.06; (2.5 mg/ml nicotine) = t(19)=4.82, p=0.0001; (10 mg/ml nicotine) = t(23)=2.22, p<0.05); (50 mg/ml nicotine) = t(24)=0.10, p=0.92). (Figure 1D). The response to nicotine vapor resembled an inverted-U, with the greatest change in preference to 2.5 mg/ml nicotine e-liquid and a decreasing preference that coincided with increasing nicotine concentrations. Specifically, there was a significant difference between the highest and lowest nicotine concentrations tested (i.e., 2.5 mg/ml and 50 mg/ml nicotine; one-way ANOVA: F(3,96)=3.08, p<0.05; Tukey’s multiple comparisons: p<0.05). Preference scores between mice conditioned with nicotine and those conditioned with vehicle did not differ (Figure 1D).

### A strawberry additive is not rewarding

We next investigated the possible influence of the strawberry e-cigarette liquid additive on CPP. We first sought to identify the major chemical components found in the commercial strawberry additive used in these studies (Liquid Barn™, Simi Valley, CA), which we expected would contain several volatile chemicals that together determine the “strawberry” odor profile. We sampled the volatiles from the strawberry additive using qualitative headspace gas chromatography/mass spectrometry (GC/MS). This analysis uncovered nine primary constituents, including ethyl butyrate, 2-methyl-ethyl butyrate, benzyl acetate, propyl butyrate, 3-hexen-1-ol, 3-hexenyl acetate, linalool, menthyl acetate, and benzyl butyrate, in order of decreasing chromatographic peak responses (Figure 1-1). No nicotine was detected in the strawberry additive sample (Table 1-1).

Before studying the possible effect of the strawberry additive on nicotine reward or aversion, we investigated its inherently rewarding or aversive properties by assaying the responses of mice to the additive-containing vapor with our CPP procedure. The vaporized strawberry additive did not produce CPP on its own regardless of concentration (2.5% or 10% of the e-liquid solution; one-sample t-tests vs 0s, not shown), and there were no differences between treatment groups (one-way ANOVA: F(2,58)=0.56, p=0.57) (Figure 1E).

**Figure 1-1.**
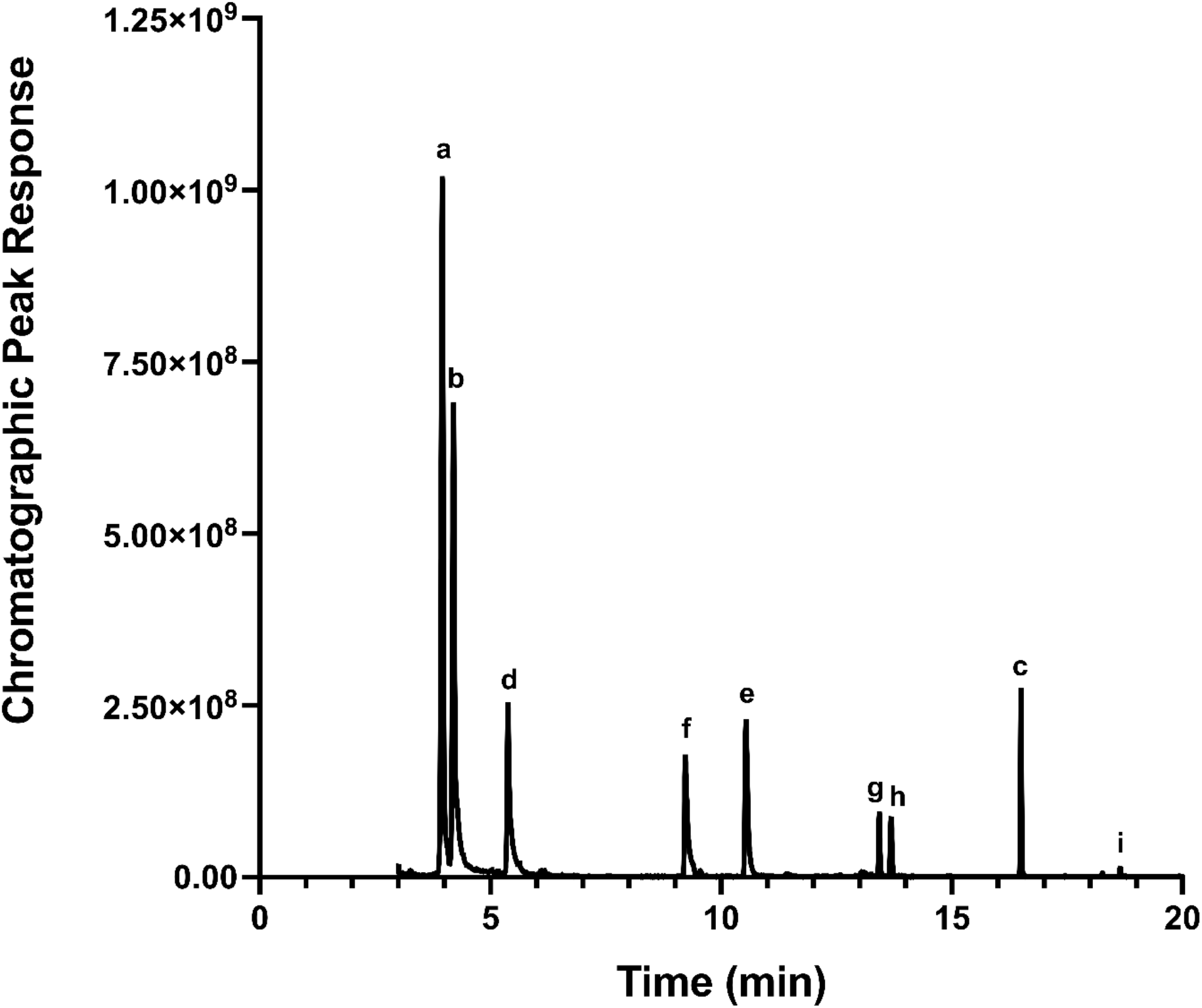
GC/MS detects chemical volatiles in strawberry e-liquid. The chromatogram from GC/MS analysis of 0.1% “Strawberry Flavor Concentrate” (Liquid Barn™) in water shows the following chemicals dominate in the commercial e-liquid headspace in order from highest to lowest chromatographic peak responses: (a) ethyl butyrate, (b) 2-methyl-ethyl butyrate, (c) benzyl acetate, (d) propyl butyrate, (e) 3-hexen-1-ol, (f) 3-hexenyl acetate, (g) linalool, (h) menthyl acetate, and (i) benzyl butyrate.

### Strawberry additive does not alter nicotine CPP

We then assessed the effect of strawberry additive on nicotine preference. Preference scores for vapor generated from nicotine e-liquid alone (first shown in Figure 1D) were compared to those for vapor containing the same concentrations of nicotine plus strawberry additive (2.5% v/v) (Figure 2). Strawberry additive did not significantly affect CPP for the nicotine-paired compartment (two-way ANOVA: F_[nicotine conc.]_(2,138)=4.80, p<0.01, F_[additive]_(1,138)=3.70, p=0.06, F_[interaction]_(2,138)=1.49, p=0.23) (Figure 2).

**Figure 2.**
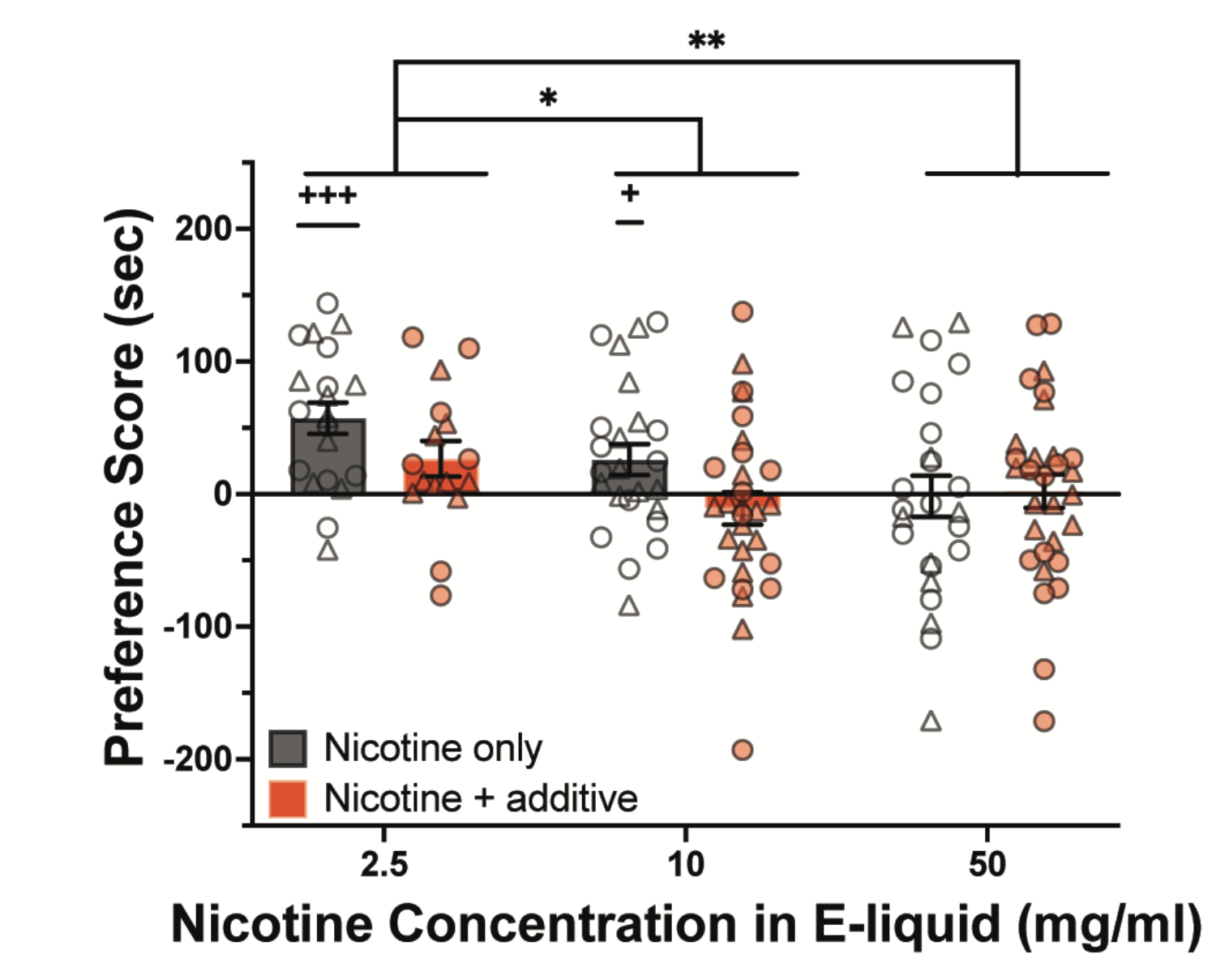
Strawberry additive does not affect nicotine CPP. Adolescent mice were conditioned to nicotine vapor with or without a strawberry additive using a CPP-biased protocol (Nicotine only: n= 20, 24, 25; Nicotine + additive: n= 16,29,30). Scatter plots represent individual mice and bars represent the mean ± sem. One sample t-tests (vs 0 second change in time spent in CS+): + p<0.05, +++ p<0.001. Two-way ANOVA and multiple comparisons: *p<0.05, **p<0.01. For all charts, circles represent individual male mice and triangles represent female mice.

### A strawberry additive in nicotine e-cigarette vapors leads to increased plasma cotinine concentrations

Despite its increased translational value, nicotine exposure via vapor is inherently more variable than via injections (due to, for example, individual differences in length and depth of inhalation), complicating attempts to standardize exposure. We therefore measured the plasma concentration of cotinine—the primary metabolite of nicotine—in mice undergoing CPP to obtain a more precise biological readout of nicotine delivery.

Surprisingly, despite the e-liquids containing the same concentration of nicotine, exposures to nicotine vapor containing a strawberry additive appeared to yield higher plasma cotinine levels at the highest nicotine concentrations (Figure 3A). This phenomenon was not apparent at the lowest nicotine concentration (2.5 mg/ml), possibly reflecting the low precision of ELISA quantification at the cotinine levels produced by this nicotine concentration (e.g., 2.0-9.7 ng/ml). Thus, the 2.5 mg/ml nicotine concentration was excluded from our analyses.

**Figure 3.**
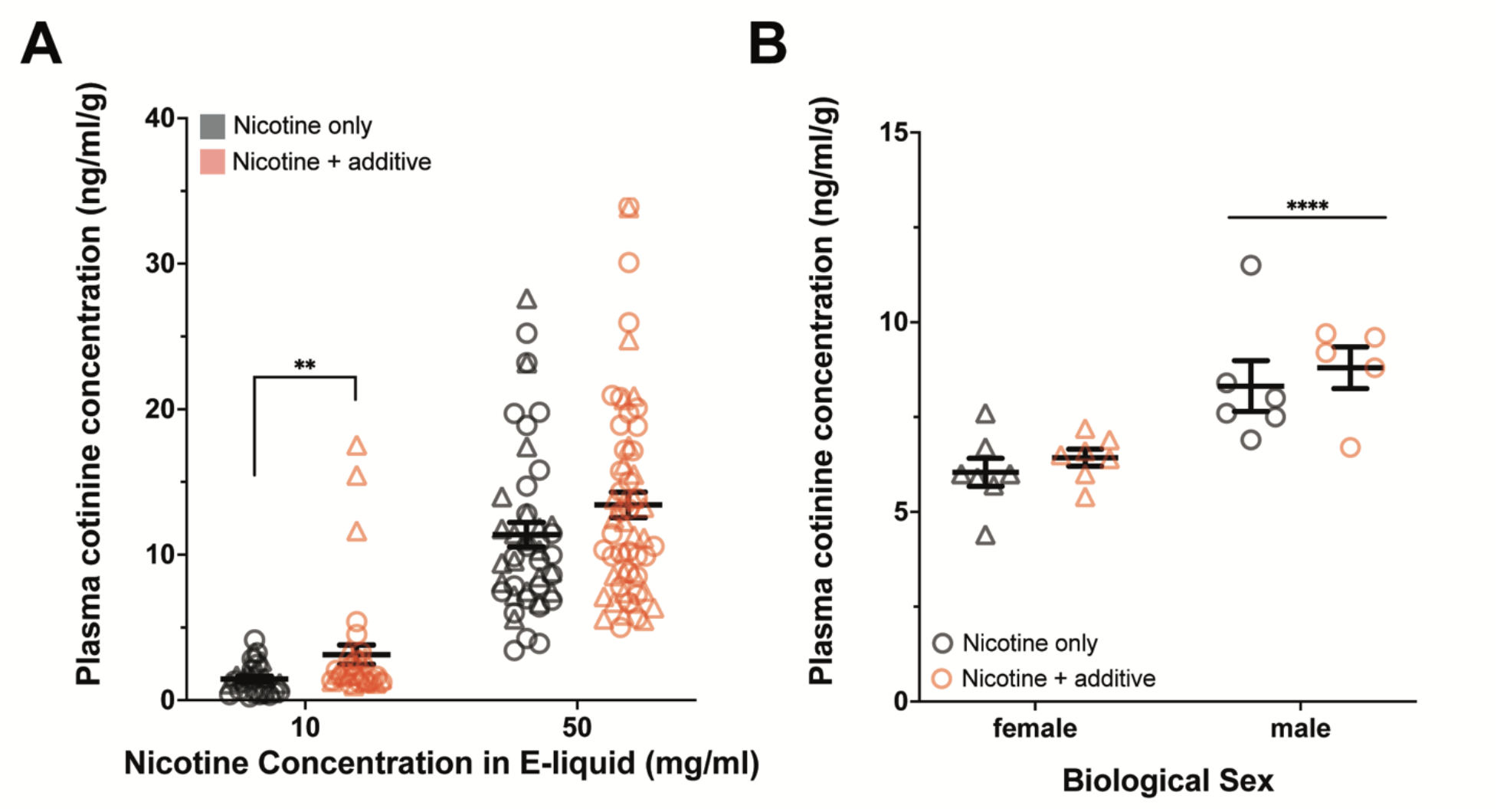
Increased plasma cotinine levels in mice exposed to strawberry nicotine vapors, a proxy for increased nicotine exposure. (A) Mice exposed to vapor containing a strawberry-additive and nicotine e-liquid (10 or 50 mg/ml) had higher plasma cotinine concentrations relative to their body weight (ng/ml/g) than mice exposed to nicotine only vapor. (10 mg/ml nicotine: n=28, +additive: 34; 50 mg/ml nicotine: n=46, +additive: 60). (B) Adolescent mice were interperitoneally injected with 1.0 mg/kg nicotine in PBS with or without 2.5% strawberry additive. The presence of strawberry additive in the injection vehicle did not influence plasma cotinine concentrations 30-minutes post-injection, however there was a significant effect of sex where, overall, male mice had higher plasma cotinine concentrations 30-minutes post-nicotine injection (females: n=7, 7; males: n=6, 5). **p<0.01, ****p<0.0001. Circles represent individual male mice and triangles represent female mice.

There was a trend for increased plasma cotinine concentrations after exposure to vapor from 10 and 50 mg/ml nicotine e-liquid that also contained the strawberry additive, and a significant interaction between nicotine concentrations and the additive (Schreier-Ray-Hare, nicotine concentration*additive: H_[nicotine conc.]_=80.23, p<0.0001; H_[additive]_=3.24, p=0.07; H_[interaction]_=44.54, p<0.0001), (Figure 3A). Mann-Whitney tests were used for further comparisons (Bonferroni-corrected alpha=0.025). Strawberry additive only had a significant effect on plasma cotinine concentrations following the 10 mg/ml nicotine treatment (Mann-Whitney tests: 10 mg/ml nicotine vs. 10 mg/ml nicotine + strawberry: p<0.01; 50 mg/ml nicotine vs. 50 mg/ml nicotine + strawberry: p= 0.11) (Figure 3A). The difference between the nicotine only and nicotine + additive groups at the 10 mg/ml nicotine concentration was “modest” (Cohen’s d=0.58), whereas the effect at the 50 mg/ml nicotine concentration was “small” (d=0.33) (Cohen, 1988).

A difference in nicotine exposure levels could hypothetically alter the behavioral response to nicotine vapor with strawberry additive independent of the additive’s sensory and rewarding properties. In other words, it is possible that the negative trend of additive on preference scores was related to increased nicotine intake (see Figure 2). To address this potential caveat, we sorted mice into “cotinine categories” based on plasma cotinine levels measured at the end of the vapor CPP experiment (see methods section “*Cotinine Category Determination”*). These “cotinine categories” allowed us to compare the behavior of mice in the additive-free and additive-containing nicotine groups that had similar levels of nicotine exposure. Interestingly, there was a negligible difference in preference scores between data categorized by concentration of nicotine in the e-liquid (as in Figure 2) and data sorted by nicotine exposure levels (i.e., “cotinine categories”; data not shown). Given this result, we present the preference and aversion scores in Figure 2 based on the concentration of nicotine in the e-liquid, consistent with the rest of the manuscript.

### Strawberry additive does not influence nicotine metabolism

We next considered the potential mechanisms underlying the effect of the strawberry additive in the e-liquid on levels of nicotine exposure (i.e. cotinine; Figure 3A). One possibility is that the additive interferes with normal nicotine metabolism and, consequently, plasma cotinine levels. To test this hypothesis, we injected adolescent mice with either 1.0 mg/kg nicotine (i.p.) in a 100% PBS vehicle or 1.0 mg/kg nicotine in a PBS solution containing 2.5% strawberry additive. There was a significant effect of sex on cotinine concentrations following nicotine injection that was not eliminated by normalizing by weight, thus sex was included as an interaction term. Inclusion of the strawberry additive did not affect plasma cotinine concentrations 30 mins post-injection (two-way ANOVA sex*additive: F_[sex]_(1,21)=25.50, p<0.0001, F_[additive]_(1,21)=0.89, p=0.36, F_[interaction]_(1,21)=0.01, p=0.92) (Figure 3B). These results suggest that the strawberry additive did not influence nicotine metabolism.

### Addition of a strawberry e-cigarette additive promotes the inhalation of nicotine vapor

We next evaluated the possibility that mice were interacting with strawberry flavored vapors differently; perhaps they inhaled more often when the e-cigarette vapor included the strawberry additive or reduced inhalation of unflavored nicotine due to its properties as an irritant (Oliveira-Maia et al., 2009). To test these hypotheses, we used a plethysmograph to measure respiration in mice while they sampled nicotine vapors (Figure 4A,B). Notably, in this design, e-cigarette liquids were not heated in an effort to more directly investigate olfactory sources of possible reward or aversion, since heating results in a more multisensory stimulus due to the sounds of the heating, visual cues of a plume, and temperature changes. Mice were first habituated to the residual sensory cues of stimuli being delivered into the plethysmograph by repeatedly delivering 50/50 VG/PG vehicle (Figure 4C) into the plethysmograph over two days. On the third day, we delivered pseudorandom trials of e-cigarette odors while simultaneously measuring sniffing (Figure 5).

**Figure 4.**
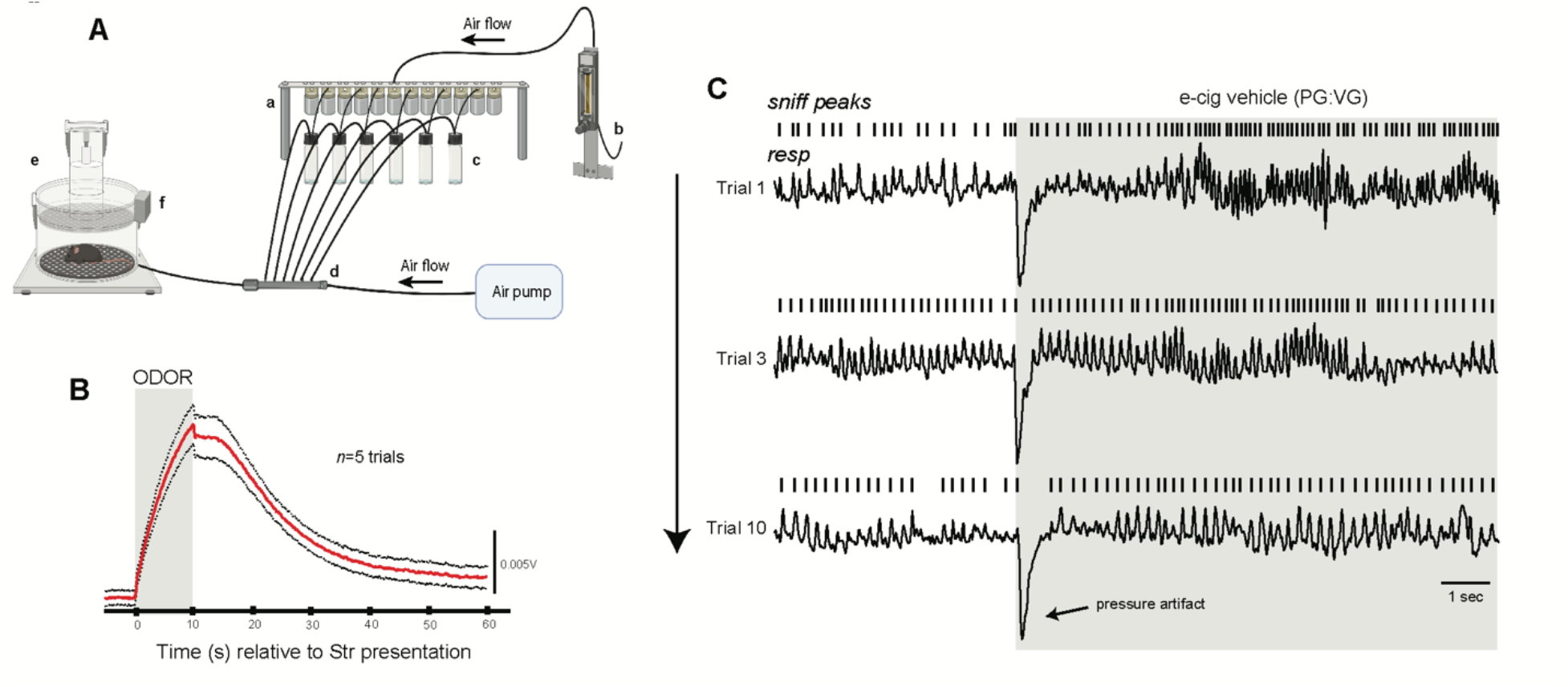
Paradigm for monitoring sniffing of mice to e-cigarette odors. (A) Design of the system. Openings of computer-controlled valves (a) allows for the flow of air as regulated by a flow meter (b) to pass through its respective valve into the connected odor vial (c). Airflow continues for 10 s, flowing from the headspace of the vial containing liquid odor through the connected tubing to ultimately combine with constant air flow in a manifold (d) and then to the mouse in the plethysmograph (e). Odor enters from the base of the plethysmograph, beneath the perforated floor. Changes in air pressure within the plethysmograph (indicating respiration) are detected by a flow transducer and digitized (f). Image made with BioRender. (B) The average of 5 trails of 10 s presentations of 2.5% strawberry (Str) e-liquid odor acquired via a photoionization detector (Aurora Scientific) illustrating onset and evacuation of odor from within the plethysmograph. Timing of strawberry odor delivery is indicated by gray box (C). Representative respiratory traces from a single mouse throughout presentations of PG/VG. Rasters indicate sniff peaks. Timing of PG/VG delivery is indicated by gray box.

**Figure 5.**
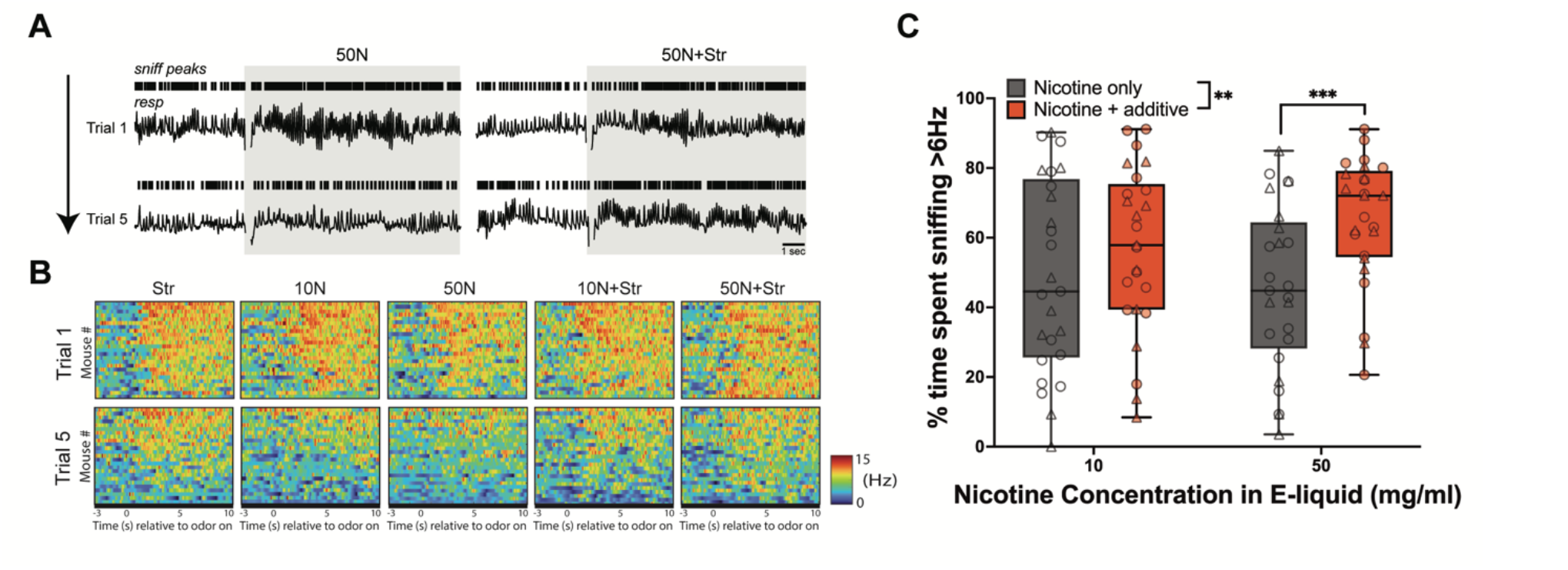
Addition of strawberry e-cig additive promotes inhalation of nicotine vapor. (A) Example respiratory traces from one mouse showing its sniffing in response to pseudorandom presentations of 50 mg/ml nicotine (50N (left)) and 50 mg/ml nicotine with the strawberry additive (50N+Str (right)). Rasters indicate sniff peaks. (B) Sniffing dynamics of all mice during the first and fifth presentation of all e-cigarette odors. 2-D histograms depict the average sniffing frequency across the 10s of odor presentation. Data are organized in descending order based on average sniff frequency during odor presentation. Dashed line indicates odor onset. (C) Percent of time sniffing by mice during presentation with flavored and unflavored nicotine vapors. Two-way ANOVA detected a main effect of flavor and multiple comparisons tests determined that mice spent significantly more time in investigatory sniffing when presented with 50N+Str. N=25, **p<0.01 and ***p<0.001 for all comparisons. In panel C, circles represent individual male mice and triangles represent female mice.

Vapors from all e-cigarette liquids elicited fast exploratory sniffing upon their first presentation (Figure 5A,B). Importantly, this indicates that the mice perceived all vapors, even nicotine when presented alone. As expected (Sundberg et al., 1982; Wesson et al., 2008), over repeated trials of odor delivery, mice habituated their sniffing response to most but not all vapors. This sustained sniffing response is meaningful in the present context since mice sniff odors they perceive as attractive more than odors they do not (*e*.*g*., (Baum and Keverne, 2002; Jagetia et al., 2018)).

We calculated the amount of time the mice spent in high-frequency investigatory sniffing [>6Hz (Wesson et al., 2008)] upon the final presentation of each vapor and found that many of the mice studied engaged in more sniffing during the final presentations of vapors containing strawberry additive alone or nicotine when it contains strawberry additive (Figure 5B). We detected a main effect of strawberry additive on investigatory sniffing towards nicotine vapor (Two-way ANOVA, nicotine*additive: F_[additive]_(1,24)=8.36, p<0.01; F_[nicotine]_(1,24)=0.50, p=0.49; F_[interaction]_(1,24)=2.97, p=0.10), with subsequent *post hoc* tests revealing that this effect was specific to the 50 mg/ml nicotine e-liquid (Sidak’s multiple comparisons test, p<0.001) (Figure 5C). There were no differences in the amplitude of sniffing between the different e-cigarette vapors (p>0.05; data not shown). These results indicate that mice actively inhale strawberry containing e-cigarette odor more than they do e-cigarette odor with nicotine alone.

## Discussion

We used a vapor delivery system to investigate how a commercial e-cigarette additive (*i*.*e*., strawberry flavor) might impact nicotine-associated behaviors. Our observations suggest that a strawberry additive may increase exposure to nicotine by increasing the interaction with nicotine vapor rather than altering reward. Our results support the hypothesis that additives with significant chemosensory attributes, specifically those fruit-like in their composition, promote nicotine intake.

### Adolescent mice express conditioned reward in response to nicotine vapor

We found that nicotine vapor leads to CPP in adolescent mice (Figure 1D). Interestingly, our results suggest that, when nicotine is delivered via inhalation, reward may occur at lower levels of exposure than previously thought. The strongest place preference to nicotine was observed at the lowest nicotine e-liquid concentration, which yielded plasma cotinine levels (≈8 ng/ml) that were most likely lower than those observed when measuring CPP in response to nicotine injection (0.3 and 0.5 mg/kg) (Fudala et al., 1985; Jorenby et al., 1990; Torres et al., 2008). These studies did not measure plasma cotinine concentrations. However, when we measured cotinine levels following intraperitoneal injections of 0.3 and 0.5 mg/kg we observed plasma cotinine concentrations of ≈46 ng/ml and ≈76 ng/ml, respectively. The level of CPP observed here is consistent with that reported in CPP experiments using systemic injections (Brielmaier et al., 2008; Brunzell et al., 2009; Sershen et al., 2010). The detection of CPP at low nicotine vapor concentrations could be due to the increased rate of nicotine delivery to the brain when inhaled rather than injected, which can affect subjective reward (Jensen et al., 2020; Samaha and Robinson, 2005). However, at least one other study has also demonstrated CPP in adolescents at a seemingly “subthreshold” dose (*i*.*e*., 0.1 mg/kg nicotine, subcutaneous injection) (Ahsan et al., 2014).

An important consideration is that mice and humans interact differently with e-cigarette vapors. In humans, e-cigarette use is highly multisensory: users inhale vapors orally (resulting in perception of taste, temperature, and retronasal olfaction) as well as nasally, which together provide important information on “flavor”. In contrast, mice are preferentially nasal breathers, so that while some exposure occurs orally, the majority occurs nasally (Agrawal et al., 2008). Trigeminal fibers are also present in the mouse nose, where they detect pungent stimuli (e.g., high concentrations of nicotine vapor), together with the olfactory receptors that detect volatile compounds, including nicotine and the chemicals found in vapor additives (e.g., Figure 1-1). Thus, although our data do not perfectly reproduce the human experience, they nevertheless provide relevant information on how fruity additives might alter the perception of nicotine vapor in humans.

### A strawberry additive does not alter the expression of nicotine vapor reward

We did not find evidence that the strawberry additive altered adolescent nicotine reward; however, methodological factors may have affected this. For example, we chose to measure the effect of a relatively low concentration of strawberry additive (2.5% v/v) based on the mouse’s sensitive olfactory system and on our finding no significant effect on preference scores at 10% (Figure 1F,G). Additives in the e-liquids used by human vapers, however, are often present at much higher concentrations—up to twice that of the nicotine present (Omaiye et al., 2019). It is possible that, at higher additive concentrations, more volatiles would be transferred into the e-cigarette vapor and absorbed, where they might have pharmacological effects capable of modifying nicotine reward (Avelar et al., 2019).

Consumer e-cigarette products emit a vast number of volatiles. Indeed, a comprehensive study of 277 commercial e-liquids identified 155 unique sensory volatiles, with an average e-liquid containing ∼25 chemicals (Omaiye et al., 2019). Our finding that a single commercial e-liquid, dominated in its volatile composition by only nine chemicals, does not alter nicotine reward or aversion should not be generalized. It is possible that other individual flavor components, such as farnesol, or even the same chemicals found in strawberry—at different concentrations or in combination with others—could alter the response to nicotine and affect nicotine CPP. Indeed, farnesol, an e-cigarette additive that mimics “green apple” flavor, can enhance nicotine reward in male mice (Avelar et al., 2019). The difference between our report and (Avelar et al., 2019) might also reflect differences in pharmacology between farnesol and the volatiles present in the strawberry additive studied here or, alternatively, could result from the different methods used. Avelar et al. injected nicotine and farnesol rather than delivering them as vapor, potentially leading to significantly higher farnesol absorption levels than those reached via inhalation. In addition, pyrolysis during vaping might generate compounds with different pharmacological properties than those of the original additive.

### Strawberry additive increases nicotine exposure in adolescent mice

Mice exposed to strawberry additive-containing nicotine vapor had higher plasma cotinine concentrations than those exposed to nicotine alone. We first considered the possibility that the strawberry additive could affect nicotine metabolism and result in increased plasma levels of cotinine, the primary metabolite of nicotine. In fact, linalool—a common volatile compound in e-cigarette flavors, which we detected as a constituent in the Liquid Barn™ additive (Figure 1-1)— affects cytochrome P450 activity in the rat liver, albeit only at very high doses (Nosková et al., 2016). However, when we injected nicotine with or without the presence of the strawberry additive, we did not observe a meaningful effect on nicotine metabolism and plasma cotinine levels, suggesting that altered metabolism is not driving a difference in plasma cotinine levels.

Additives may also alter nicotine absorption through changes in pH of e-liquid solutions and vapors (Patten and De Biasi, 2020; St.Helen et al., 2018, 2017), as has been previously reported in humans (St.Helen et al., 2018, 2017). Nicotine absorption is facilitated by more acidic vapors, which allow a larger proportion of the nicotine to reach the lung where it can be buffered, absorbed, and rapidly delivered to the brain (Patten and De Biasi, 2020). Although we did not explicitly test for pH effects, we observed a trend for a slightly lower pH in nicotine e-liquids containing the strawberry flavorant (data not shown). Thus, it is possible that more acidic strawberry e-liquids/vapors lead to an increase in nicotine absorption, contributing to higher plasma cotinine concentrations (Figure 3A).

Although altered chemical properties of e-cigarette vapors may have contributed to higher nicotine exposure, our study suggests that chemosensory-rich additives found in e-cigarette liquids alter the way the mice interact with e-cigarette vapors. Additives appear to influence vaping topography in humans (Audrain-McGovern et al., 2016; St.Helen et al., 2018); St. Helen and colleagues measured a longer average puff duration when participants vaped a strawberry e-liquid compared to a tobacco e-liquid (St.Helen et al., 2018). This phenomenon is recapitulated by our results demonstrating increased time spent sniffing vapors containing the strawberry additive, which may contribute to the increased nicotine intake we observed. The observed effect size in our study ranges from relatively small to modest, nevertheless the odorant’s inclusion increased the level of plasma cotinine between 114% (10 mg/ml) and 18% (50 mg/ml). Given the popularity of flavored e-cigarette liquids, even a seemingly small difference may have a profound impact at a population level, and a small increase in nicotine exposure may be sufficient to increase susceptibility or rate of developing dependence (MacPherson et al., 2008; DiFranza et al., 2007), including within days or weeks of inhaling their first cigarette (DiFranza et al., 2007). Detecting early symptoms of nicotine dependence is important because it may help identify individuals on trajectories toward heavier tobacco use (Lessov-Schlaggar et al., 2008).

Our preclinical results add to the argument that additives with significant chemosensory attributes, specifically those fruit-like in their composition, influence the intake of e-cigarette vapors in manners that promote nicotine intake. Because mice are preferential nasal breathers (Agrawal et al., 2008), we predict the olfactory components of the additive greatly mediate this effect. However, additional work is needed to resolve the sensory influences of this effect.

## Conclusion

Our results indicate that a strawberry fruit additive, one of many in a category of highly used “e-cigarette flavors”, significantly increases nicotine exposure but not reward. This finding is particularly important for understanding e-cigarette effects on adolescents who are vulnerable to nicotine uptake and whose brains are sensitive to long-term disruptions mediated by nicotine exposure at this age (DiFranza et al., 2007; Goriounova and Mansvelder, 2012; Substance Abuse and Mental Health Services Administration, 2011; Yuan et al., 2015). Our data also highlight an underappreciated role of e-cigarette flavor additives: their ability to alter one’s response to nicotine via sensory contributions, possibly leading to differences in nicotine exposure. Further research using models of nicotine self-administration with a fruit- or sweet-flavored vapor may help predict the ultimate impact of increased nicotine intake under these conditions.

## References

Agrawal A, Singh SK, Singh VP, Murphy E, Parikh I (2008) Partitioning of nasal and pulmonary resistance changes during noninvasive plethysmography in mice. J Appl Physiol 105:1975–1979.

Ahsan HM, de la Peña JBI, Botanas CJ, Kim HJ, Yu GY, Cheong JH, Peña JBI de la, Botanas CJ, Kim HJ, Yu GY, Cheong JH (2014) Conditioned place preference and self-administration induced by nicotine in adolescent and adult rats. Biomol Ther (Seoul) 22:460–466.

Audrain-McGovern J, Strasser AA, Wileyto EP (2016) The impact of flavoring on the rewarding and reinforcing value of e-cigarettes with nicotine among young adult smokers. Drug Alcohol Depend 166:263–267.

Avelar AJ, Akers AT, Baumgard ZJ, Cooper SY, Casinelli GP, Henderson BJ (2019) Why flavored vape products may be attractive: Green apple tobacco flavor elicits reward-related behavior, upregulates nAChRs on VTA dopamine neurons, and alters midbrain dopamine and GABA neuron function. Neuropharmacology 158:107729.

Bacon WE, Snyder HL, Hulse SH (1962) Saccharine preference in satiated and deprived rats. J Comp Physiol Psychol 55:112–114.

Baum M, Keverne E (2002) Sex difference in attraction thresholds for volatile odors from male and estrous female mouse urine. Horm Behav 41:213–219.

Brielmaier JM, McDonald CG, Smith RF (2008) Nicotine place preference in a biased conditioned place preference design. Pharmacol Biochem Behav 89:94–100.

Brunzell DH, Mineur YS, Neve RL, Picciotto MR (2009) Nucleus Accumbens CREB Activity is Necessary for Nicotine Conditioned Place Preference. Neuropsychopharmacology 2009 34:8 34:1993–2001.

Centers for Disease Control and Prevention (2019) Progress Erased: Youth Tobacco Use Increased During 2017-2018 [WWW Document]. URL https://www.cdc.gov/media/releases/2019/p0211-youth-tobacco-use-increased.html (accessed 1.22.20).

Chaffee BW, Watkins SL, Glantz SA (2018) Electronic cigarette use and progression from experimentation to established smoking. Pediatrics 141.

Chen JC, Green K, Fryer C, Borzekowski D (2019) Perceptions about e-cigarette flavors: a qualitative investigation of young adult cigarette smokers who use e-cigarettes. Addiction Research & Theory 27:420–428.

Cohen J (1988) Statistical Power Analysis for the Behavioral Sciences Second Edition, 2nd ed. Lawrence Erlbaum Associates.

Cooper SY, Akers AT, Henderson BJ (2021) Flavors Enhance Nicotine Vapor Self-administration in Male Mice. Nicotine & Tobacco Research 23:566–572.

Creamer MR, Wang TW, Babb S, Cullen KA, Day H, Willis G, Jamal A, Neff L (2019) Tobacco Product Use and Cessation Indicators Among Adults — United States, 2018. MMWR Morb Mortal Wkly Rep 68:1013–1019.

Desor JA, Beauchamp GK (1987) Longitudinal changes in sweet preferences in humans. Physiol Behav 39:639–641.

DiFranza JR, Savageau JA, Fletcher K, Pbert L, O’Loughlin J, McNeill AD, Ockene JK, Friedman K, Hazelton J, Wood C, Dussault G, Wellman RJ (2007) Susceptibility to nicotine dependence: The development and assessment of nicotine dependence in youth 2 study. Pediatrics 120:e974–e983.

Dufour VL, Arnold MB (1966) Taste of saccharin as sufficient reward for performance. Psychol Rep 19:1293–4.

Ford A, MacKintosh AM, Bauld L, Moodie C, Hastings G (2016) Adolescents’ responses to the promotion and flavouring of e-cigarettes. Int J Public Health 61:215–224.

Fudala P, Teoh K, Iwamoto E (1985) Pharmacologic characterization of nicotine-induced conditioned place preference. Pharmacol Biochem Behav 22:237–241.

Gadziola M, Tylicki K, Christian D, Wesson D (2015) The olfactory tubercle encodes odor valence in behaving mice. J Neurosci 35:4515–4527.

Garrett PI, Honeycutt SC, Marston C, Allen N, Barraza AG, Dewey M, Turner B, Peterson AM, Hillhouse TM (2021) Nicotine-free vapor inhalation produces behavioral disruptions and anxiety-like behaviors in mice: Effects of puff duration, session length, sex, and flavor. Pharmacol Biochem Behav 206:173207.

Goldenson NI, Kirkpatrick MG, Barrington-Trimis JL, Pang RD, McBeth JF, Pentz MA, Samet JM, Leventhal AM (2016) Effects of sweet flavorings and nicotine on the appeal and sensory properties of e-cigarettes among young adult vapers: Application of a novel methodology. Drug Alcohol Depend 168:176–180.

Goriounova NA, Mansvelder HD (2012) Short- and Long-Term Consequences of Nicotine Exposure during Adolescence for Prefrontal Cortex Neuronal Network Function. Cold Spring Harb Perspect Med 2:a012120.

Hoffman AC, Salgado RV, Dresler C, Faller RW, Bartlett C (2016) Flavour preferences in youth versus adults: a review. Tob Control 25:ii32–ii39.

Jagetia S, Milton A, Stetzik L, Liu S, Pai K, Arakawa K, Mandairon N, Wesson DW (2018) Inter- and intra-mouse variability in odor preferences revealed in an olfactory multiple-choice test. Behavioral neuroscience 132:88–98.

Jensen KP, Valentine G, Gueorguieva R, Sofuoglu M, KP J, G V, R G, M S (2020) Differential effects of nicotine delivery rate on subjective drug effects, urges to smoke, heart rate and blood pressure in tobacco smokers. Psychopharmacology (Berl) 237:1359–1369.

Johnson ME, Bergkvist L, Mercado G, Stetzik L, Meyerdirk L, Wolfrum E, Madaj Z, Brundin P, Wesson DW (2020) Deficits in olfactory sensitivity in a mouse model of Parkinson’s disease revealed by plethysmography of odor-evoked sniffing. Scientific Reports 2020 10:1 10:1–13.

Jorenby DE, Steinpreis RE, Sherman JE, Baker TB (1990) Aversion instead of preference learning indicated by nicotine place conditioning in rats. Psychopharmacology 1990 101:4 101:533–538.

Kim H, Lim J, Buehler SS, Brinkman MC, Johnson NM, Wilson L, Cross KS, Clark PI (2016) Role of sweet and other flavours in liking and disliking of electronic cigarettes. Tob Control 25:ii55–ii61.

Leventhal A, Cho J, Barrington-Trimis J, Pang R, Schiff S, Kirkpatrick M (2019) Sensory attributes of e-cigarette flavours and nicotine as mediators of interproduct differences in appeal among young adults. Tob Control.

Leventhal AM, Goldenson NI, Barrington-Trimis JL, Pang RD, Kirkpatrick MG (2019) Effects of non-tobacco flavors and nicotine on e-cigarette product appeal among young adult never, former, and current smokers. Drug Alcohol Depend 203:99–106.

Levin ED, Rose JE, Behm F (1990) Development of a citric acid aerosol as a smoking cessation aid. Drug Alcohol Depend 25:273–279.

Li Q, Zhan Y, Wang L, Leischow SJ, Zeng DD (2016) Analysis of symptoms and their potential associations with e-liquids’ components: A social media study. BMC Public Health 16:1–12.

McKendrick G, Graziane NM (2020) Drug-Induced Conditioned Place Preference and Its Practical Use in Substance Use Disorder Research. Front Behav Neurosci 14:173.

Mennella JA, Bobowski NK, Reed DR (2016) The development of sweet taste: From biology to hedonics. Rev Endocr Metab Disord 17:171–178.

Moline JM, Golden AL, Highland JH, Wilmarth KR, Kao AS (2000) Health effects evaluation of theatrical smoke, haze, and pyrotechnics.

Montanari C, Kelley LK, Kerr TM, Cole M, Gilpin NW (2020) Nicotine e-cigarette vapor inhalation effects on nicotine & cotinine plasma levels and somatic withdrawal signs in adult male Wistar rats. Psychopharmacology (Berl) 237.

Nosková K, Dovrtělová G, Zendulka O, Řemínek R, Juřica J (2016) The Effect of (-)-Linalool on the Metabolic Activity of Liver CYP Enzymes in Rats. Physiol Res 65:499–504.

Oliveira-Maia AJ, Stapleton-Kotloski JR, Lyall V, Phan T-HTHT, Mummalaneni S, Melone P, Desimone JA, Nicolelis MALL, Simon SA (2009) Nicotine activates TRPM5-dependent and independent taste pathways. Proc Natl Acad Sci U S A 106.

Omaiye EE, McWhirter KJ, Luo W, Tierney PA, Pankow JF, Talbot P (2019) High concentrations of flavor chemicals are present in electronic cigarette refill fluids. Sci Rep 9:2468.

Patten T, De Biasi M (2020) History repeats itself: Role of characterizing flavors on nicotine use and abuse. Neuropharmacology 177:108162.

Pepper JK, Ribisl KM, Brewer NT (2016) Adolescents’ interest in trying flavoured e-cigarettes. Tob Control 25:ii62–ii66.

Rose JE, Zinser MC, Tashkin DP, Newcomb R, Ertle A (1984) Subjective response to cigarette smoking following airway anesthetization. Addictive behaviors 9:211–5.

Samaha AN, Robinson TE (2005) Why does the rapid delivery of drugs to the brain promote addiction? Trends Pharmacol Sci 26:82–87.

Sershen H, Hashim A, Lajtha A (2010) Differences between nicotine and cocaine-induced conditioned place preferences. Brain Res Bull 81:120–124.

Siu EC, Tyndale RF (2007) Characterization and comparison of nicotine and cotinine metabolism in vitro and in vivo in DBA/2 and C57BL/6 mice. Mol Pharmacol 71:826–834.

Soneji S, Barrington-Trimis JL, Wills TA, Leventhal AM, Unger JB, Gibson LA, Yang J, Primack BA, Andrews JA, Miech RA, Spindle TR, Dick DM, Eissenberg T, Hornik RC, Dang R, Sargent JD (2017) Association Between Initial Use of e-Cigarettes and Subsequent Cigarette Smoking Among Adolescents and Young Adults. JAMA Pediatr 171:788–797.

St.Helen G, Dempsey DA, Havel CM, Jacob P, Benowitz NL (2017) Impact of e-liquid flavors on nicotine intake and pharmacology of e-cigarettes. Drug Alcohol Depend 178:391–398.

St.Helen G, Shahid M, Chu S, Benowitz NL (2018) Impact of e-liquid flavors on e-cigarette vaping behavior. Drug Alcohol Depend 189:42–48.

Substance Abuse and Mental Health Services Administration (2011) Results from the 2010 National Survey on Drug Use and Health: Summary of National Findings, NSDUH Series H-41.

Sundberg H, Døving K, Novikov S, Ursin H (1982) A method for studying responses and habituation to odors in rats. Behav Neural Biol 34:113–119.

Torres O v, Tejeda HA, Natividad LA, O’Dell LE (2008) Enhanced vulnerability to the rewarding effects of nicotine during the adolescent period of development. Pharmacol Biochem Behav 90:658–663.

Varughese S, Teschke K, Brauer M, Chow Y, Van Netten C, Kennedy SM (2005) Effects of theatrical smokes and fogs on respiratory health in the entertainment industry. Am J Ind Med 47:411–418.

Wesson DW, Donahou TN, Johnson MO, Wachowiak M (2008) Sniffing behavior of mice during performance in odor-guided tasks. Chem Senses 33:581–596.

Wieslander G, Norbäck D, Lindgren T (2001) Experimental exposure to propylene glycol mist in aviation emergency training: acute ocular and respiratory effects. Occup Environ Med 58:649.

Yuan M, Cross SJ, Loughlin SE, Leslie FM (2015) Nicotine and the adolescent brain. Journal of Physiology 593:3397–3412.

Zandstra EH, de Graaf C (1998) Sensory perception and pleasantness of orange beverages from childhood to old age. Food Qual Prefer 9:5–12.

