## Supplemental Figure 1-1 for "Strawberry additive increases nicotine vapor sampling and systemic exposure but does not enhance Pavlovian-based nicotine reward in mice"

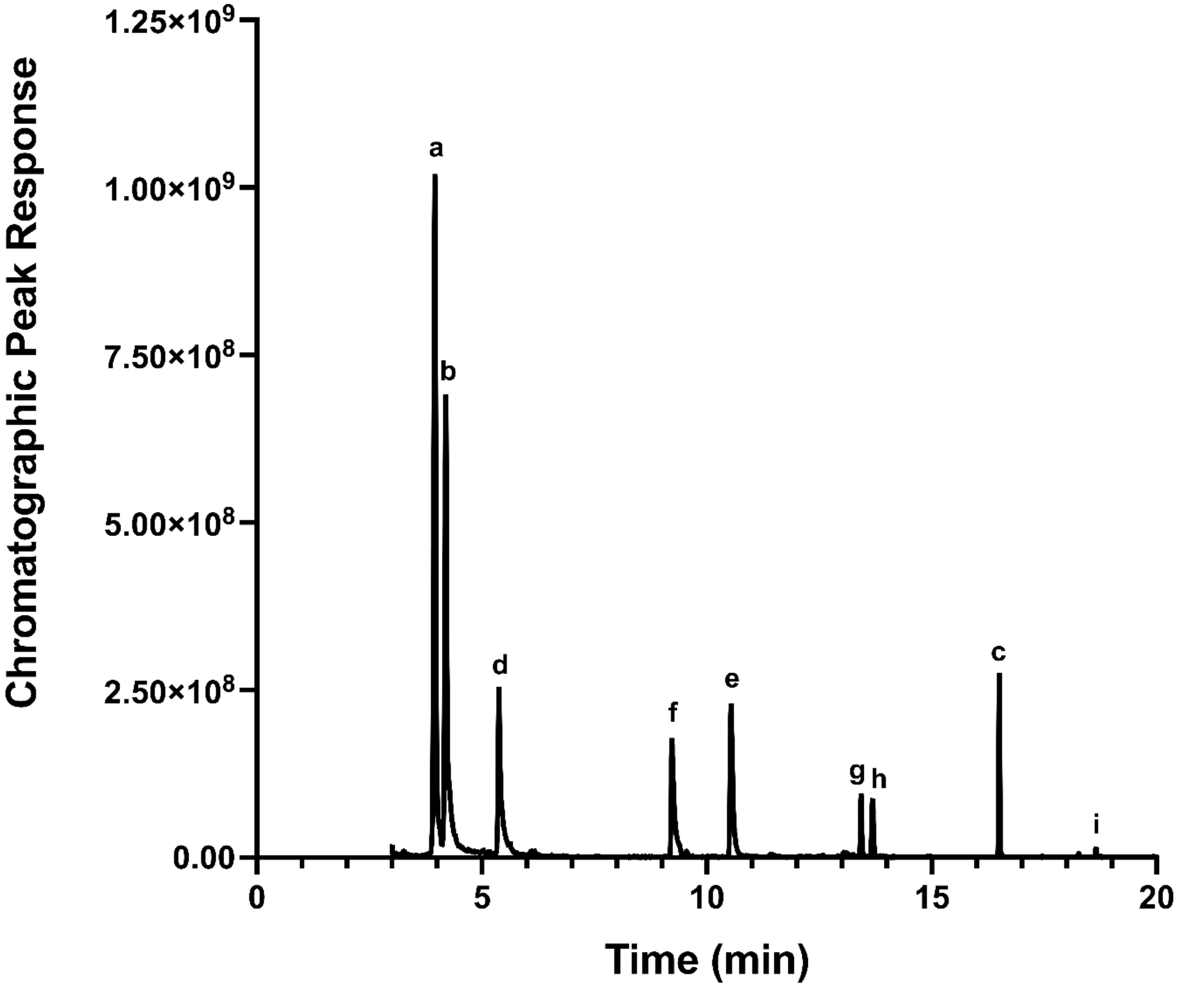


**Figure 1-1.GC/MS detects chemical volatiles in strawberry e-liquid.** The chromatogram from GC/MS analysis of 0.1% “Strawberry Flavor Concentrate” (Liquid Barn™) in water shows the following chemicals dominate in the commercial e-liquid headspace in order from highest to lowest chromatographic peak responses: (a) ethyl butyrate, (b) 2-methyl-ethyl butyrate, (c) benzyl acetate, (d) propyl butyrate, (e) 3-hexen-1-ol, (f) 3-hexenyl acetate, (g) linalool, (h) menthyl acetate, and (i) benzyl butyrate.
